# Information Retrieval using Machine Learning for Biomarker Curation in the Exposome-Explorer

**DOI:** 10.1101/2020.12.20.423685

**Authors:** Andre Lamurias, Sofia Jesus, Vanessa Neveu, Reza M Salek, Francisco M Couto

## Abstract

In 2016, the International Agency for Research on Cancer, part of the World Health Organization, released the Exposome-Explorer, the first database dedicated to biomarkers of exposure for environmental risk factors for diseases. The database contents resulted from a manual literature search that yielded over 8500 citations, but only a small fraction of these publications were used in the final database. Manually curating a database is time-consuming and requires domain expertise to gather relevant data scattered throughout millions of articles. This work proposes a supervised machine learning approach to assist the previous manual literature retrieval process.

The manually retrieved corpus of scientific publications used in the Exposome-Explorer was used as training and testing sets for the machine learning models (classifiers). Several parameters and algorithms were evaluated to predict an article’s relevance based on different datasets made of titles, abstracts and metadata.

The top performance classifier was built with the Logistic Regression algorithm using the title and abstract set, achieving an F2-score of 70.1%. Furthermore, from 705 articles classified as relevant, we extracted 545 biomarkers, including 460 new candidate entries to the Exposome-Explorer database.

Our methodology reduced the number of articles to be manually screened by the database curators by nearly 90%, while only misclassifying 22.1% of the relevant articles. We expect that this methodology can also be applied to similar biomarkers datasets or be adapted to assist the manual curation process of similar chemical or disease databases.

## Background

Biomarkers are biological parameters objectively measured in the body as indicators of normal biological conditions, environmental lifestyles, pathological conditions, or responses to therapeutic interventions [1]. They can be chemicals, metabolites, enzymes or other biochemical substances, like products of an interaction between a compound and a target molecule or a cell type. Characterizing the relationship between biomarkers and the possible biological outcomes is crucial to correctly predict clinical responses, screen, monitor and diagnose patients and to improve efficiency in clinical trials. Biomarkers play a significant role in risk assessment, as they allow one to identify exposure to hazards and to associate responses with the probability of a disease or exposure outcome.

Biomarkers of exposure are a specific type of biomarkers that reflect exposure of an individual to an environmental factor (such as diet, pollutants or infectious agents) known to affect the etiology of diseases. Compounds can get in contact with living organisms through absorption, inhalation or ingestion and then are either metabolized, stored or eliminated. This exposure can be detected by analysing biospecimens, such as blood or urine, or by measuring concentrations and characterizing the exogenous substance, its metabolites or its products of interaction with target molecules.

Exposome-Explorer is a manually curated database of biomarkers of exposure to environmental risk factors developed by the International Agency for Research on Cancer and with collaboration with the University of Alberta, Canada [2]. It contains detailed information on the nature of biomarkers, populations and subjects in which biomarkers have been measured, samples analysed, methods used for biomarker analysis, their concentrations in biospecimens, correlations with external exposure measurements, and biological reproducibility over time [3]. To develop the database, experts searched biomedical literature in the Web of Science (WOS), using queries with specific keywords associated with dietary and pollutant biomarkers. More than 8500 citations from 1975 to 2014 were manually screened according to the title and abstract to identify publications describing biomarkers of exposure. Only a small fraction was considered relevant and data from 480 publications was manually analysed and annotated, and finally included in the database.

Gathering relevant data scattered throughout millions of articles from text repositories is a time-consuming task which requires specialized professionals to manually retrieve and annotate relevant information within the articles. Keeping biological databases updated as new papers are released, as well as collecting new data, is equally challenging and time consuming. Such tasks would benefit from being assisted with text-mining tools. To our knowledge, there is no machine learning Information Retrieval (IR) solution available to assist literature screening regarding biomarkers of exposure.

Studies have been carried out to either improve the IR task using machine learning or to perform entity recognition (ER) and information extraction (IE) on biomarkers’ data. However, none applies IR-based methods to biomarkers of exposure. Almeida et al. [4] developed a machine learning system for supporting the first task of the biological literature manual curation process, called triage, which involves identifying very few relevant documents among a much larger set of documents. They were looking for articles related to characterized lignocellulose-active proteins of fungal origin to curate the mycoCLAP database [5]. They compared the performance of various classification models, by experimenting with dataset sampling factors and a set of features, as well as three different machine learning algorithms (Naïve Bayes, Support Vector Machine and Logistic Model Trees). The most fitting model to perform text classification on abstracts from PubMed was obtained using domain relevant features, an under-sampling technique, and the Logistic Model Trees algorithm, with a corresponding F-measure of 0.575. Lever et al. [6] used a supervised learning approach to develop an IE-based method to extract sentences containing relevant relationships involving biomarkers from PubMed abstracts and Pubmed Central Open Access full text papers. With this approach, they built the CIViCmine knowledge base [7], containing over 90,992 biomarkers associated with genes, drugs and cancer types. Their goal was to reduce the time needed to manually curate databases, such as the Clinical Interpretation of Variants in Cancer (CIViC) knowledgebase [8], and to make it easier for the community curators, as well as editors, to contribute with content.

Following the previous approaches, this work aims to reduce the time, effort and resources necessary to keep the Exposome-Explorer database updated as new articles are published, by using a supervised machine learning approach to automatically classify relevant publications and automatically recognize candidate biomarkers to be reviewed by the curators. The approach of the curators of this database consisted in developing search queries to retrieve relevant publications and then manually analyse each one. However, the number of publications retrieved is still too large to manually screen each one of them. This work proposes a system, available on Github (https://github.com/lasigeBioTM/BLiR), to further narrow down the literature that holds important information about biomarkers of exposure. The existing manually curated data used to develop the Exposome-Explorer database has been used to train and test the models (classifiers). We also provide a corpus articles classified by our system as relevant, along with biomarkers automatically annotated on the abstracts of these articles. When given a new publication, these classifiers can predict whether this publication is relevant to the database and annotate the candidate biomakers mentioned on that document.

## Methods

### Exposome-Explorer dataset

This work was developed using the data used to set up and develop the Exposome-Explorer, which included:

– the queries used to search for citations with information about dietary and pollutant biomarkers in the Web of Science (WOS);
– the WOS search results based on the previous queries, with 8575 citations used to manually screen the relevant articles containing biomarker information;
– the 480 publications used to extract information about biomarkers for the database.

### Data Collection

All 480 publications used to curate the database were expected to be listed in the 8575 citations retrieved from the WOS. However, only 396 of them were present: the 84 publications absent from the WOS query results were additionally identified by database annotators while screening the literature for relevant articles. These 84 scientific papers were excluded from the dataset used to build the models since we could not replicate the original workflow if we included them.

The existing dataset, listed above, was missing some features that we wanted to explore to construct our models, such as number of citations and PubMed ID. For this reason, PubMed was used to extract the titles, abstracts and metadata (publication date, author names, number of times the article was cited and journal name). The PubMed search and retrieval of PMIDs, titles, abstract and metadata was carried out with E-utilities, a public API available at NCBI Entrez system. Some publications were found through the DOI to PMID (PubMed ID) converter and others by a combined search with the title and first author name. The resulting corpus of articles consisted of 7083 publications.

### Data Preprocessing

After retrieving the title, abstract and metadata for each article, it was necessary to prepare the textual data to be used as input by the machine learning models (classifiers). This task included:

i. *Assign labels to each article:* A supervised learning approach was used to build the classifiers, which means each article (document) has a known class assigned to it. To label each article, the list with the 396 articles used to curate the database was cross-referenced with the 7083 publications in the corpus. If they were present in the list, they were considered relevant and assigned the label 1. If they were not present in the list, as they were not used to extract information about biomarkers, they were considered irrelevant and therefore assigned the label 0;
ii. *Natural Language Processing:* The text was separated into words (tokens). All words with the same root were reduced to a common form (stemming) and all stop words were removed. The tokens were lowercased and then combined into n-grams, a contiguous sequence of *n* items from a given sample of text or speech. For example, for *n* = 2, the features from the sentence *“Determining thiocyanate serum levels”*, were combined into three n-grams: *“determin thiocyan”, “thiocyan serum”* and *“serum level”*;
iii. *Transform textual data to numerical data:* The machine learning model expects numeric data as its input. However, the titles, abstracts, and metadata are in text format. To this end, each distinct token that occurred in the documents was mapped to a numerical identifier, and the number was used to refer to the token instead of the token itself;
iv. *Build the matrices:* Each feature represents one column and each document represents a row of the matrix. Depending on the type of matrix chosen, the matrix contained either n-gram counts (number of times each term occurs in each document) or TFIDF (term frequency-inverse document frequency) features (how important a n-gram is to a document in a collection of documents). An additional column was added to the training and testing data, with the respective labels. The goal of the classifier was to predict this column when applied to a new data.

The metadata of each article was handled slightly differently from the titles and the abstracts. Since it already had numerical attributes (publication date and number of citations), the matrix was created with two columns dedicated to these features, instead of having one column for each year and number of citations. The authors’ names were joined into one single word (Wallace RB *→* WallaceRB) and were neither combined into n-grams nor went through the stemming and stop word removal stages. The journal name had no special preprocessing.

Stemming was performed using the class `SnowballStemmer` from the module `nltk.stem` in the NLTK package [9]. Steps (ii), (iii) and (iv) used the Scikit-learn [10] classes `CountVectorizer` and `TfidfVectorizer` from the module `sklearn.feature extraction.text`. The main difference between the two classes is that the first one converts a collection of raw text documents to a matrix of token counts and the last one to a matrix of TFIDF features. Combinations of three different parameters were tested to preprocess the data, resulting in different matrices used to build the classifier and, therefore, different results. The parameters tested were:

– `ngram_range (min n, max n)`: the lower and upper boundary of n for different n-grams to be extracted. The range values tested were *n* = {1}, *n* = {1, 2} and *n* = {1, 2, 3};
– `min_df`: ignore all n-grams that have a document frequency lower than the given threshold. If `min_df` = 2, then terms that only appear in one article (document) will be ignored. The values of `min_df` ranged from 2 to 23, depending on the value of n used in the `ngram_range` parameter ([1 + *n − gram*, 21 + *n − gram*]);
– type of the matrix: matrix of token counts or TFIDF features.

Finally, we divided the dataset into a train set of 70% and a test set of 30%, while keeping the same proportion of positive and negative classes on both subsets. The train set was used to optimize the parameters through Cross Validation (CV) and the test set was used to obtain the results on held-out data.

### Machine learning models

The goal of the IR task was to reduce the time needed to screen the articles, by narrowing down the literature available to a set of publications that provide a reliable resource of information, in this specific case, related to biomarkers of exposure. Thus, in this case we can model the IR task as a classification task, where we have to decide whether a document is relevant or not.

#### Building the classifiers

The machine learning models, also known as classifiers, were separately trained and tested using the titles, abstracts, titles + abstracts and titles + metadata, to assess which section of the article was more suitable to predict its relevance. We explored the combination of titles and metadata since our preliminary results indicated that the metadata by itself would not obtain reasonable results. However, these preliminary results also indicated that combining the abstracts with metadata would result in equal or worse results than using just the abstracts. For this reason, we did not explore the option of combining abstracts with metadata, or combining all three.

Six machine learning algorithms were explored:

- **Decision Tree**[11]: the features are fractioned in branches that represent a condition to be applied to each instance;
- **Logistic Regression**[12]: learns a logistic function to perform binary classification;
- **Naïve Bayes**[13]: the independence of the features is assumed and a probability model is used to determine the most probable label for each instance;
- **Neural Network**[14]: this algorithm can learn non-linear functions by introducing hidden layers between the input features and output label;
- **Random Forest**[15]: combines various tree estimators trained on sub-samples of the training data;
- **Support Vector Machine**[16]: the data is represented as points in a hyperplane and the algorithm tries to establish a clear division between the instances with the same label.

The Scikit-learn package was used to run these algorithms. Most of the parameters used for each algorithm were the default ones, however, a few ones were altered to better suit the data (`class_weight, solver, kernel, gamma, bootstrap, n estimators`), others to maximize the performance of the model (`C, alpha, max_depth, min_samples_leaf`), and one to assure a deterministic behaviour during fitting (`random_state`). The values of the parameters altered to maximize the performance of the model were found through grid search with 10-fold CV on the train set. Table 1 summarizes the Scikit-learn functions used and the parameters changed for each algorithm.

**Table 1:**
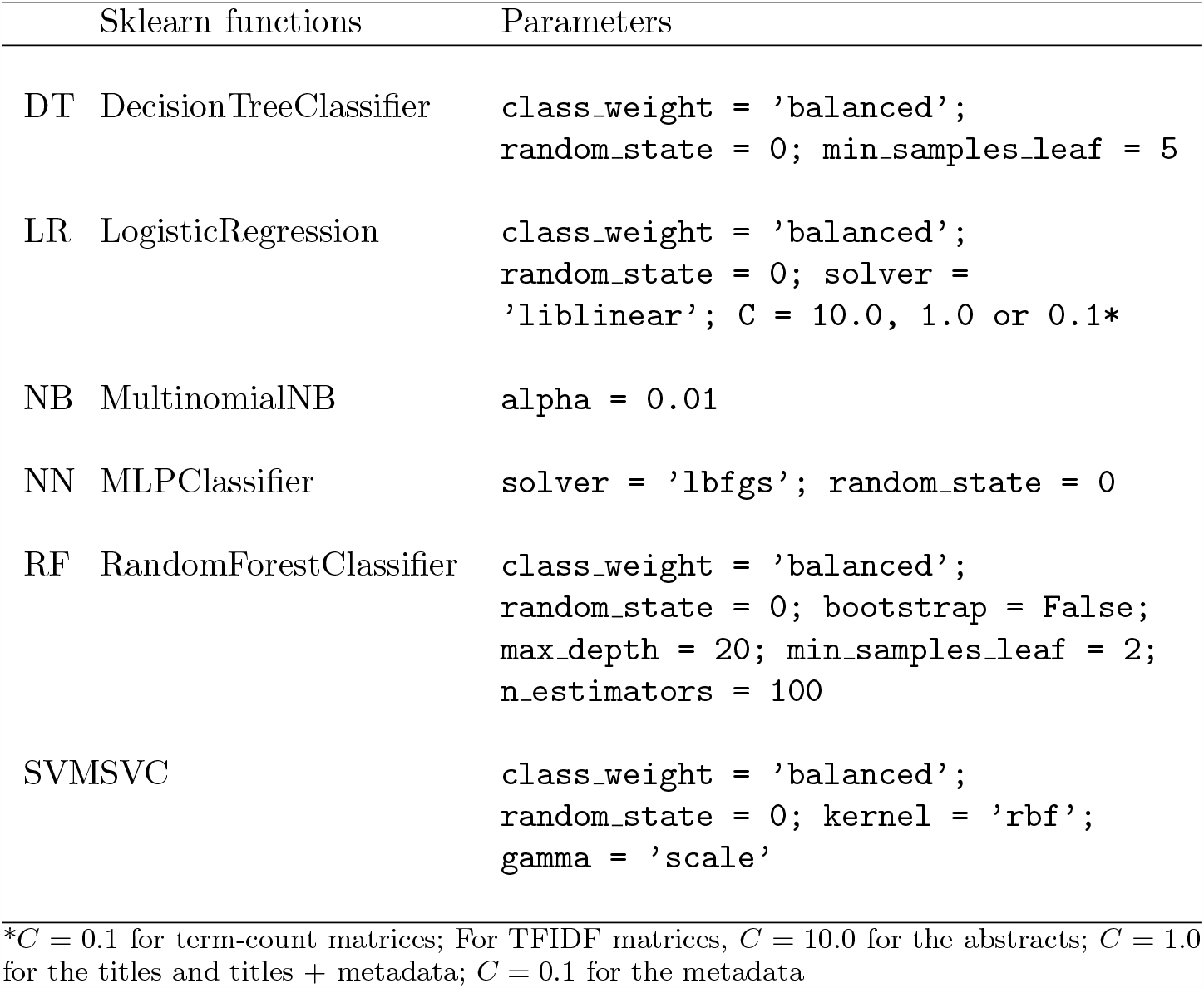
Scikit-learn functions and parameters for each algorithm: Decision Tree (DT), Logistic Regression (LR), Naïve Bayes (NB), Neural Network (NN), Random Forest (RF) and Support Vector Machine (SVM).

#### Ensemble learning

When testing different classifiers using the abstracts, titles, titles + abstracts or the titles + metadata set, the prediction each model makes for a certain article might differ. The titles + metadata model may correctly identify a publication as being relevant, while the abstracts model fails to do so. For this reason, we explored ensembles of classifiers to understand if we could retrieve more relevant publications this way.

We used two ensemble approaches to join the results of multiple models. The first was Bagging, where the same algorithm is used to train a classifier on random subsets of the training data, and the results are then combined [17]. The second was Stacking, which consists in training multiple classifiers and using their output to train a final model which predicts the classes [18]. With this approach, each of the first-level classifiers can be specified, as well as the final classifier. Therefore, we used all of the previously mentioned algorithms as first-level classifiers, and then tried each of them as the final estimator. For the Bagging approach, we also tried every algorithm previously mentioned. In both cases, we used the parameters specified in Table 1, using the Scikit-learn implementations and the default parameters of the `BaggingClassifier` and `StackingClassifier` classes.

### Performance Evaluation

In the data preprocessing task, labels were given to each article: 0 for irrelevant (negative) and 1 for relevant (positive). These labels were considered the gold-standard and represent the actual class of the publications.

In the document classification task, all classifiers built were optimized using the Scikit-learn CV function (`sklearn.model_selection.cross_validate)`. This model optimization technique provides a more accurate estimate of the model’s performance, since it evaluates how the model will perform on unseen data. Additionally, we selected a test set to evaluate the models after parameter optimization.

The `cv` parameter of the function determines how many groups the data will be split into. In this work, a *cv* = 10 was used, which means the data was separated into 10 groups, each one was used 9 times as a training set and once as the testing set. Ten different models were built using the same parameters, with different training sets. Each time a trained model was applied to testing data, it generated a vector with predicted classes for those documents. By comparing the predictions of the testing set to the gold standard, it was possible to separate the documents into four different categories:

– True Positives (TP): documents correctly labelled as positive;
– False Positives (FP): documents incorrectly labelled as positive;
– True Negatives (TN): documents correctly labelled as negative;
– False Negatives (FN): documents incorrectly labelled as negative.

This categorization allows to calculate the precision and recall, two commonly used metrics that assess the performance of the tools by measuring the quality of the results in terms of relevancy. Precision (P) is the proportion of true positives items over all the items the system has labelled as positive. Recall (R) is the proportion of true positives items over all the items that should have been labelled as positive.

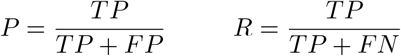

The F1-score is a measure between 0 and 1 that combines both precision and recall. Higher values of this metric indicate that the system classifies most items in the correct category, therefore having low numbers of FP and FN.

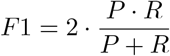

Furthermore, we also considered a variation of the F1-score, the F2-score, where more weight is given to the recall:

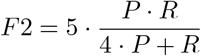

This metric was important for our evaluation since we wanted to avoid low recall values, which would mean that many documents were mistakenly classified as not relevant. Our objective was to reduce the number of documents that manual curators had to analyse, but without losing important information.

To estimate the balance between the true positive rate (recall) and false positive rate, we also computed the area under the Receiver operating characteristic (ROC AUC). We used the Scikit-learn implementation of this measure, which computes the area under a curve plotted by the true positive rate and false positive rate at various thresholds.

### Biomarkers recognition

We performed biomarker recognition on the documents classified as positive by our best performing classifier. The objective of this task was to demonstrate how the document classifiers can be used to aid the curation process. By automatically screening the articles for biomarkers, curators can focus on articles that mention entities of their interest and help them to extract information from those articles. We used the Chemical Entities of Biological Interest (ChEBI) ontology [19] as a reference vocabulary, and identified chemical entities mentioned in the abstracts using MER, a Minimal Entity Recognition tool [20]. This tool returns a list of entities recognized in the text, including their exact location and unique identifier, if available. It has the advantage of being adaptable to any lexicon and has been tested in a scenario where thousands of documents had to be processed [21]. Unlike other entity recognition tools, MER does not require training data to recognize a new type of entity. In this case, the entities correspond to candidate biomarkers, along with documents where they were found.

## Results

### Data Collection and Preprocessing

After data collection, the Exposome-Explorer dataset consisted of titles, abstracts, and metadata from a total of 7083 publications. Among them, 6687 were considered irrelevant, because no information about biomarkers was extracted from them for the Exposome-Explorer database. The remaining 396 publications were considered relevant, as they were used to construct the database.

In the beginning, all articles from all types of biomarkers in the dataset were used, however, this approach yielded poor results. To try to improve the results, the data was restricted to articles regarding dietary biomarkers, since they were handled more attentively by the curators. The new dataset consisted of 3016 publications (2860 irrelevant + 156 relevant).

### Document Classification

#### Dietary biomarker publications

Our first objective was to train models to classify which articles from a search query were relevant to the Exposome-Explorer database. We optimized both the parameters used to preprocess the diet training data (ngram, minimum frequency, vectorizer), as well as hyperparameters of each algorithm, using grid search-CV. For each algorithm, we tested several combinations and selected the trained models that achieved the highest score of each metric on the CV evaluation.

The maximum values each algorithm could reach for these metrics, using optimized preprocessing and algorithm parameters, are summarized in Table 2. The complete values for each highest metric, as well as the parameters used, can be found in Additional File 1. For example, the maximum F2-score of 0.701 of the LR algorithm on the titles+abstracts set was obtained using a `min_df` of 5, `ngram_range` (1, 3) and a token count matrix. We can see that all algorithms except Decision Trees could achieve high values on the various data subsets, although using only the titles, the LR algorithm achieved higher scores in most metrics. The parameters and algorithms used to maximize the F2-score for each feature set can be found in Table 4.

**Table 2:** Dietary biomarkers document classification results. Highest precision, recall, F1-score, F2-score and ROC-AUC achieved by each algorithm: Decision Tree (DT), Logistic Regression (LR), Naïve Bayes (NB), Neural Network (NN), Random Forest (RF) and Support Vector Machine (SVM). The highest value of each metric on each feature type is bolded.

| TITLES |  |  |  |  |  |
| --- | --- | --- | --- | --- | --- |
|  | MaxPrecision | MaxRecall | MaxF1 | MaxF2 | MaxROC-AUC |
| DT | 0.216 | 0.433 | 0.262 | 0.302 | 0.635 |
| LR | 0.388 | **0.707** | **0.495** | **0.601** | **0.910** |
| NB | 0.528 | 0.652 | 0.475 | 0.561 | 0.903 |
| NN | 0.560 | 0.331 | 0.385 | 0.348 | 0.887 |
| RF | 0.415 | 0.661 | 0.489 | 0.577 | 0.889 |
| SVM | **0.586** | 0.688 | 0.462 | 0.560 | 0.904 |

| ABSTRACTS |  |  |  |  |  |
| --- | --- | --- | --- | --- | --- |
|  | MaxPrecision | MaxRecall | MaxF1 | MaxF2 | MaxROC-AUC |
| DT | 0.337 | 0.570 | 0.397 | 0.449 | 0.720 |
| LR | 0.559 | 0.770 | 0.618 | **0.687** | 0.953 |
| NB | 0.606 | 0.752 | **0.644** | 0.684 | 0.952 |
| NN | **0.854** | 0.441 | 0.542 | 0.472 | 0.948 |
| RF | 0.621 | 0.705 | 0.591 | 0.644 | **0.954** |
| SVM | 0.512 | **0.798** | 0.572 | 0.658 | 0.949 |

| TITLES + ABSTRACTS |  |  |  |  |  |
| --- | --- | --- | --- | --- | --- |
|  | MaxPrecision | MaxRecall | MaxF1 | MaxF2 | MaxROC-AUC |
| DT | 0.379 | 0.533 | 0.419 | 0.480 | 0.740 |
| LR | 0.550 | 0.779 | 0.614 | **0.701** | 0.948 |
| NB | 0.601 | 0.788 | **0.643** | 0.695 | 0.950 |
| NN | **0.764** | 0.432 | 0.528 | 0.463 | 0.945 |
| RF | 0.665 | 0.687 | 0.586 | 0.634 | **0.953** |
| SVM | 0.512 | **0.807** | 0.564 | 0.664 | 0.948 |

| TITLES + METADATA |  |  |  |  |  |
| --- | --- | --- | --- | --- | --- |
|  | MaxPrecision | MaxRecall | MaxF1 | MaxF2 | MaxROC-AUC |
| DT | 0.307 | 0.468 | 0.328 | 0.390 | 0.693 |
| LR | 0.392 | **0.734** | **0.502** | **0.616** | **0.930** |
| NB | 0.373 | 0.707 | 0.456 | 0.566 | 0.914 |
| NN | **0.603** | 0.266 | 0.333 | 0.274 | 0.905 |
| RF | 0.476 | 0.725 | 0.492 | 0.592 | 0.924 |
| SVM | 0.163 | 0.168 | 0.144 | 0.154 | 0.725 |

In addition to exploring single classifiers, we also explored two ensemble approaches: Bagging and Stacking. We trained a Stacked classifier that combined the best individual models (Table 1), and then applied again one of the algorithms as the final classifier. Table 3 show the maximum Precision, Recall, F1-Score, F2-score and ROC-AUC of each algorithm, using the Stacking and Bagging approach, and training only on the abstracts+titles subset, which provided the best results of most of the individual models. This way, we can compare directly with the results of Table 2. The full set of values of each metric is also provided in Additional File 1.

**Table 3:** Dietary biomarkers ensemble classifiers’ results. Highest precision, recall, F2-score and ROC-AUC reached for each algorithm: Decision Tree (DT), Logistic Regression (LR), Naïve Bayes (NB), Neural Network (NN), Random Forest (RF) and Support Vector Machine (SVM). The NB algorithm did not work with the Stacking approach.

| BAGGING |  |  |  |  |  |
| --- | --- | --- | --- | --- | --- |
|  | MaxPrecision | MaxRecall | MaxF1 | MaxF2 | MaxROC-AUC |
| DT | **0.742** | 0.477 | 0.546 | 0.502 | 0.941 |
| LR | 0.664 | **0.707** | **0.642** | **0.663** | 0.950 |
| NB | 0.716 | 0.580 | 0.594 | 0.582 | 0.951 |
| NN | 0.729 | 0.477 | 0.537 | 0.497 | 0.942 |
| RF | 0.648 | 0.541 | 0.571 | 0.550 | **0.958** |
| SVM | 0.740 | 0.405 | 0.489 | 0.433 | 0.957 |

| STACKING |  |  |  |  |  |
| --- | --- | --- | --- | --- | --- |
|  | MaxPrecision | MaxRecall | MaxF1 | MaxF2 | MaxROC-AUC |
| DT | 0.417 | 0.735 | 0.513 | 0.622 | 0.842 |
| LR | 0.380 | **0.890** | 0.521 | **0.685** | **0.961** |
| NN | **0.581** | 0.505 | 0.521 | 0.509 | 0.911 |
| RF | 0.568 | 0.734 | **0.624** | 0.673 | 0.952 |
| SVM | 0.374 | **0.890** | 0.513 | 0.679 | 0.947 |

**Table 4:** Algorithm and parameters used to get the highest F2 for each set of data.

|  | Title | Abstracts | T + A | T + M |
| --- | --- | --- | --- | --- |
| Algorithm | LR | LR | LR | LR |
| df | 4 | 4 | 5 | 4 |
| n-gram | [1, 2] | [1, 3] | [1, 2] | 2 |
| matrix | token-count | TFIDF | token-count | token-count |
| Precision | 0.388 | 0.500 | 0.514 | 0.392 |
| Recall | 0.707 | 0.770 | 0.779 | 0.734 |
| F1-score | 0.495 | 0.587 | 0.614 | 0.502 |
| F2-score | 0.601 | 0.687 | 0.701 | 0.616 |
| ROC AUC | 0.910 | 0.938 | 0.937 | 0.930 |

We then applied the classifiers of the previous table with highest F2-score to the test set which we did not use during grid search-CV 5. With this held-out dataset, we wanted to observe if the classifiers had been overfitted to the training set due to the parameter optimization procedure.

#### All biomarker publications

To quantify how much restricting the dataset to dietary biomarkers had improved the results, new models were trained with the whole corpus of 7083 publications from all biomarkers using the same algorithms and parameters that had maximized the recall score for dietary biomarkers. The comparison between the values of precision, recall and F-score can be found in Table 6.

Afterwards, we performed biomarker recognition on the 707 articles identified by the classifier with highest F2 on Table 6 as relevant (which included TPs and FPs), using an automatic annotation tool and manual validation. We obtained 545 biomarkers from these documents, which we provide as Additional File 2.

## Discussion

The highest F2-score (0.701) was obtained using a single classifier with the Logistic Regression (LR) algorithm on the abstracts and title set, using cross-validation (Table 2). Among the 905 dietary publications used to test the classifiers, 365 were classified as positive, which could reduce by 90% (proportion of articles classified as positive) the time needed to find 77.9% (recall score) of the relevant articles, and only 22.1% of the relevant articles would be lost. Looking at the results from the titles and metadata set, globally lower values were obtained when compared to the abstracts sets. Using features from both the titles and abstracts resulted in better F2-scores in almost every algorithm, comparing with using them separately. This indicates that, similarly to how it is carried out during manual curation, both titles and abstracts should be considered when evaluating the relevance of an article to the database. The LR algorithm obtained the best performance on many metrics, although the SVM algorithm obtained a higher recall using the titles and titles and abstracts, and Random Forests obtained the highest ROC-AUC on the same sets. The Neural Networks algorithm obtained the highest precision using the abstracts, titles and abstracts, and titles and metadata sets.

To assess whether joining the best models would improve the scores, we applied two ensemble approaches to the abstracts set: Bagging and Stacked. In some cases, using a Bagging approach results in better scores than just the model by itself, for example, comparing the scores of the Decision Tree classifier. However, in most cases, using just one classifier provided better results. The Stacking approach also obtained better scores in some cases, including a maximum recall of 0.890 using the Logistic Regression and SVM classifiers. However this approach took much longer to train since it requires training one model with each of the previously mentioned algorithms, as well as an additional model to predict the class based on the other models’ prediction scores. Furthermore, both ensemble approaches resulted in similar or worse results than the single classifiers. This could be due to the increased complexity of these models, which may be less adaptable to new data due to overfitting to the train data.

In Table 5, we can see the effect of the cross-validation evaluation when compared to the test set validation. Although some of the scores are lower, the LR algorithm also achieves the highest balanced scores and the Neural Networks achieves the highest precision. The Stacking algorithm achieves a high recall, but at the cost of lower precision. Although the balanced metrics are lower on the test set when compared to the test set evaluation, we believe that the difference is not relevant, since the cross-validation results were averaged over five iterations, and the test set shows the results of only one run.

**Table 5:** Dietary biomarkers classifiers on the test set. Precision, recall, F2-score and ROC-AUC achieved each algorithm: Decision Tree (DT), Logistic Regression (LR), Naïve Bayes (NB), Neural Network (NN), Random Forest (RF) and Support Vector Machine (SVM), as well as the Bagging and Stacking approaches, using the combinations that achieved the highest F2-score.

|  | Precision | Recall | F1 | F2 | ROC-AUC |
| --- | --- | --- | --- | --- | --- |
| DT | 0.403 | 0.532 | 0.459 | 0.500 | 0.744 |
| LR | 0.530 | 0.745 | **0.619** | **0.689** | 0.854 |
| NB | 0.515 | 0.745 | 0.609 | 0.684 | 0.853 |
| NN | **0.700** | 0.447 | 0.545 | 0.482 | 0.718 |
| RF | 0.450 | 0.766 | 0.567 | 0.672 | 0.857 |
| SVM | 0.451 | 0.787 | 0.574 | 0.685 | 0.867 |
| Bagging | 0.542 | 0.681 | 0.604 | 0.648 | 0.825 |
| Stacking | 0.388 | **0.851** | 0.533 | 0.687 | **0.889** |

### Error analysis

In order to interpret the gap of results between the training set and the predictions obtained from the classifiers, the LR classifier built with the titles was analysed. This classifier had a similar recall score to the abstracts but, as the titles are shorter, they make the interpretation easier. One interesting pattern we noticed was that almost all titles that had the words “food frequency questionnaire” were classified as relevant. From a total of 82 titles containing these words, only 2 were classified as irrelevant (both had words such as “calcium”, “water” and “energy” that were mostly found on irrelevant articles); 29 were TP and the remaining 51 were being wrongly labelled as relevant.

The title “Toenail selenium as an indicator of selenium intake among middle-aged men in an area with low soil selenium” was classified as negative, when it was in fact used in the database (FN). 39 out of 40 titles with the word “selenium” were not used in the database and thus labelled irrelevant: this over-represented feature may be the reason why the classifier failed to classify this article as relevant although selenium was considered of interest by the annotators.

It is also important to highlight that papers inserted in the database have been analysed considering the full-texts. This means that papers tagged as “relevant” either by the classifier and/or manually, could subsequently be rejected by the annotators, for a variety of reasons including “the paper is not-available online”, or “the data in the paper is not presented in a way acceptable for the database”. These papers would then be considered false positive by the classifier, because they are present in the corpus of citations but absent from the database.

Restricting the analysis to the dietary biomarker citations provided much better metrics than when using all the data from the database (dietary, pollutants, and reproducibility values) (Table 6). When restricting the analysis to citations describing the different classes of biomarkers of pollution, the performance of the models was even lower (preliminary results not shown). This difference in performance could be explained by the difference of nature of the data searched by the annotators for the different sets of biomarkers. For dietary biomarkers, the focus was made on publications providing correlation values between dietary intakes and biomarkers measured in human biospecimens, and mostly describing validation studies of dietary questionnaires with biomarkers. For the pollutant biomarkers, the focus was made on papers describing concentration values of pollutant biomarkers in human biospecimens. Moreover, by lack of time and human resources, not all potentially relevant publications on pollution biomarkers were inserted in the Exposome-Explorer, while the dietary biomarkers were handled more attentively. As a consequence, the dietary biomarkers account for almost half of the entries of the database. All of this could explain why the model seems to perform better for dietary biomarkers. Having a closer look at false positives obtained by the classifier on pollutants could be a good way to check if the model developed on dietary biomarkers could also be applied to pollutants, and identify new relevant papers from the corpus of pollutants. This also means that as we obtain a more comprehensive corpus for other classes of biomarkers, the performance of our machine learning solution will also improve.

**Table 6:** Comparison of precision, recall and F-score between the whole dataset and the restricted dataset.

| TITLES |  |  |  |  |  |
| --- | --- | --- | --- | --- | --- |
|  | Precision | Recall | F1 | F2 | ROC AUC |
| All biomarkers | 0.362 | 0.529 | 0.430 | 0.485 | 0.737 |
| Dietary biomarkers | 0.386 | 0.574 | 0.462 | 0.523 | 0.762 |
| ABSTRACTS |  |  |  |  |  |
|  | Precision | Recall | F1 | F2 | ROC AUC |
| All biomarkers | 0.218 | 0.765 | 0.340 | 0.510 | 0.801 |
| Dietary biomarkers | 0.342 | 0.809 | 0.481 | 0.635 | 0.862 |
| TITLES + ABSTRACTS |  |  |  |  |  |
|  | Precision | Recall | F1 | F2 | ROC AUC |
| All biomarkers | 0.354 | 0.630 | 0.453 | 0.545 | 0.781 |
| Dietary biomarkers | 0.468 | 0.787 | 0.587 | 0.693 | 0.869 |
| TITLES + METADATA |  |  |  |  |  |
|  | Precision | Recall | F1 | F2 | ROC AUC |
| All biomarkers | 0.310 | 0.681 | 0.426 | 0.550 | 0.795 |
| Dietary biomarkers | 0.330 | 0.638 | 0.435 | 0.538 | 0.784 |

### Biomarkers recognition

We manually confirmed the biomarkers detected and provide the full list as an Additional File. This list contains 545 chemical entities, along with the documents where each one was identified. The most mentioned entities were “polychlorobiphenyl” (CHEBI:53156), “acid” (CHEBI:37527) and “phthalate” (CHEBI:17563). While some of these entities already existed on the Explorer-Explorer database, most (460) did not match any existing entry. In some cases, the results of this approach were to broad, for example, “acid” corresponds to mentions to specific acid molecules such as acetic acid or uric acid. However, this approach was able to identify biomarkers on abstracts that were ignored during the development of the database. For example, our approach identified “omega-6 fatty acid” as a candidate biomarker (CHEBI:36009) on a document that was not used for the database. This list of chemicals and documents where they were found can be used to generate new candidate entries for the Exposome-Explorer database.

## Conclusions

The Exposome-Explorer database is being manually curated, without any assistance from machine learning tools. As the number of scientific papers continues to grow, text-mining tools could be a great help to assist the triage of documents containing information about biomarkers of exposure and keep the database updated.

To this end, several machine learning models were created using different combinations of preprocessing parameters and algorithms. These classifiers were trained using the publications’ abstracts, titles and metadata. The model with the highest F2-score (70.1%) was built with the LR algorithm and used the titles and abstracts to predict a paper’s relevance. We extracted named-entities from the abstracts selected by this model, obtaining a total of 545 candidate biomarkers.

To apply this methodology to the database curation pipeline, the IR task will consist of two steps. In the first one, articles will be retrieved using the query search on WOS, to target domain-specific publications. Then, the classifier could be used to narrow down the publications even more, and a named-entity recognition tool can be used to provide candidate entries to the database. Manual curation will still be needed, to extract information about biomarkers from full-text articles.

In the future, we will work on improving the results from the classifiers that use the metadata set. For example, by testing different weights to the authors, according to the position they appear in, or by creating new features that result from the combinations between all authors within the same article. We will also study the impact of the recognized biomarkers in the retrieval classification. When analysing why the model misclassified some publications, a few chemicals, like “calcium” and “selenium” were strongly associated with irrelevant articles. An idea to explore is to replace chemical tokens by a category they belong, such as “chemical”, and see if it improves the precision and recall of the classifiers. This should avoid over-fitting on the training set. Furthermore, we will improve this gold standard by adding more types of biomarkers, that can also classify non-dietary biomarkers. Another idea to explore is to train deep learning models on other biomedical corpora and apply to the documents of this gold standard dataset, since these approaches require larger datasets.

## Abbreviations

WOS: Web of Science;
IR: Information Retrieval;
ER: Entity Recognition
IE: Information Extraction;
CIViC: Clinical Interpretation of Variants in Cancer;
CV: Cross Validation
TFIDF: Term Frequency–Inverse Document Frequency;
TP: True Positives;
FP: False Positives;
TN: True Negatives;
FN: False Negatives;
ChEBI: Chemical Entities of Biological Interest;
PAH: Polycyclic aromatic hydrocarbons;
PCB: Polychlorinated biphenyls;
HCA: Heterocyclic amines;
PCDD/F: Polychlorodibenzo-p-dioxins and Polychlorodibenzo-furans;
PBDE: Polybrominated diphenyl ethers;
PBB: Polybrominated biphenyls.

## Ethics approval and consent to participate

Not applicable.

## Consent for publication

Not applicable.

## Availability of the data and material

The code and data used is available at https://github.com/lasigeBioTM/BLiR.

## Author’s contributions

All authors read and approved the final manuscript. Where authors are identified as personnel of the International Agency for Research on Cancer / World Health Organization, the authors alone are responsible for the views expressed in this article and they do not necessarily represent the decisions, policy or views of the International Agency for Research on Cancer / World Health Organization.

## Additional Files

**Additional file 1**

Excel file with the precision, recall, F1-score, F2-score and ROC-AUC of the classifiers for all combinations of preprocessing parameters (min df, ngram range and matrix type) and algorithms (Decision Tree, Logistic Regression, Naïve Bayes, Neural Network, Random Forest and SVM), along with two ensemble algorithms, Bagging and Stacking.

**Additional file 2 — Candidate Biomarkers**

Tab-separated values file with the ChEBI IDs of the biomarker entities found on the abstracts classified as relevant, along with the PubMed IDs of the documents where they were found.

